# iTRAQ-Based Proteomic Profiling of PLC-Mediated Expression of 20E-Induced Protein in *Apolygus lucorum* (Meyer-Dür)

**DOI:** 10.1101/2020.10.15.341115

**Authors:** Yong-An Tan, Xu-Dong Zhao, Jing Zhao, Qin-Qin Ji, Liu-Bin Xiao, De-Jun Hao

**Affiliations:** Institute of Plant Protection, Jiangsu Academy of Agricultural Sciences, Nanjing, China, 210014; College of Forestry, Nanjing Forestry University, Nanjing, China, 210037; Taizhou Customs of the People’s Republic of China,Taizhou, China, 225300

**Keywords:** *Apolygus lucorum*, 20-hydroxyecdysone, phospholipase C, iTRAQ, Proteomics, Western-blot

## Abstract

The polyphagous pest *Apolygus lucorum* has become the dominant insect in *Bacillus thuringiensis* (Bt) cotton fields. The hormone 20-hydroxyecdysone (20E) regulates multiple events in insect development and physiology. 20E responses are controlled by pathways triggered by phospholipase C (PLC)-associated proteins. However, 20E-modulated genes whose expression is affected by PLC remain unknown. Here, isobaric tag for relative and absolute quantitation (iTRAQ) and immunoblot were carried out for comparing differentially expressed proteins (DEPs) in *A. lucorum* in response to 20E and the PLC inhibitor U73122, respectively. Totally 1624 DEPs were, respectively, found in the 20E/control, U73122/control, and 20E+U73122/control groups. Venn diagram analysis further revealed 8 DEPs that were shared among the three groups. Immunoblot validated these findings, which corroborated and highlighted the reliability of proteomics. KEGG enrichment analysis showed that the DEPs were included in diverse signaling pathways. The largest portion of DEPs among the three groups were categorized in metabolic pathways. In addition, DEPs among the three groups were also found to regulate the Ras-MAPK and PI3K-AKT pathways. This is the first time that iTRAQ was carried out to assess proteome alteration in *A. lucorum* nymphs in response to 20E and a PLC inhibitor. These findings provide novel insights into protein expression in *A. lucorum* in response to 20E, and a more comprehensive understanding of the function of PLC in 20E signal transduction.

## 1. INTRODUCTION

*Apolygus lucorum* (Meyer-Dür), a worldwide agricultural pest, has become a key insect pest in cotton fields (Lu and Wu, 2008). With the rising plantation area of transgenic Bt cotton cultivars, *Helicoverpa armigera* (Hübner) has been controlled effectively (Wu *et al.*, 2005; Lu and Wu, 2011). However, *A. lucorum* is hard to control due to elevated mobility, broad host range and complex feeding properties (Lu *et al.*, 2010). At present, chemical control with different insecticides remains the most applied option for the management of *A. lucorum*. Extensive use of insecticides has resulted in *A. lucorum* resistance to these products, with inherent environmental risks (Lu *et al.*, 2007). Consequently, new tools, particularly those that target insect functions via molecular mechanisms for pest management, are urgently needed.

The insect steroid ecdysone and its active metabolite 20-hydroxyecdysone (20E) trigger and coordinate multiple events in insect development and physiology such as caste determination, diapause and longevity. Many effects of 20E are mediated by a heterodimer comprised of the ecdysone receptor (EcR) and ultraspiracle (USP) (Nakagawa and Henrich, 2009; Christiaens *et al.*, 2010). In detail, EcR contains a basic ligand-independent activation region near the N terminus, the main dimerization domain and a ligand-binding domain, which heterodimerizes with USP (Iwema *et al.*, 2007; Xu *et al.*, 2020). This pocket with remarkable flexibility can functionally interact with the molting hormone 20E. The 20E-EcR-USP complex binds to a specific DNA response element (EcRE) to regulate the transcription of target genes (Boulanger and Dura, 2015).

As soluble compounds, steroid hormones easily diffuse into the cell nucleus via the cytoplasmic membrane, and interact with nuclear steroid hormone receptors to substantially modulate gene expression (Thompson *et al.*, 1995). Meanwhile, steroid hormones trigger non-genomic signaling via the cytoplasmic membrane (Meldrum, 2007; Sandén *et al.*, 2011; Falkenstein *et al.*, 2000). Phospholipase C (PLC) constitutes a major class of regulatory enzymes contributing to non-genomic membrane pathways controlling 20E signaling genes, mostly by hydrolyzing the polar head groups in inositol-containing membrane phospholipids (Liu *et al.*, 2014; Cai *et al.*, 2014). For instance, apoptosis in silkworm anterior silk glands is induced by 20E-associated GPCR-PLC-IP_3_-Ca^2+^-PKC signaling (Iga *et al.*, 2007; Manaboon *et al.*, 2009). In *H. armigera*, through GPCR-PLC-Ca^2+^ signaling, 20E induces a rapid phosphorylation of cyclin-dependent kinase 10 (CDK10) to promote gene transcription (Liu *et al.*, 2014). PLC substantially affects larva development and pupation, and 20E could increase PLC amounts during molting and metamorphosis. Via ErGPCR, Ga_q_ and Src kinases, 20E readily triggers tyrosine phosphorylation in the SH2 domain of PLC and its migration toward the cytoplasmic membrane. Through PLC/Ca^2+^ signaling, 20E induces EcRE’s transcriptional activity by modulating PKC phosphorylation at Ser 21, which is important for its interaction with EcRE (Liu *et al.*,2014; Jing *et al.*, 2015). In a previous study, we found 20E regulates AlTre-1 through AlPLCγ and alters Vg expression and ovary development, thereby facilitating female reproduction in *A. lucorum* (Tan *et al.*, 2020). These data indicated that PLC participates in 20E signal transduction in a non-genomic pathway.

Currently, differential proteomics can help identify proteins with physiological significance. Recently, isobaric tags for relative and absolute quantitation (iTRAQ) were designed for the simultaneous assessment of 4 to 8 distinct specimens by MS/MS, thereby increasing throughput and reducing assay-related errors (Ross *et al.*, 2004; Ye *et al.*, 2010). Additionally, this novel method is sensitive, automated and multi-dimensional (Caubet *et al.*, 2010). Here, iTRAQ was utilized to investigate proteome alterations in *A. lucorum* in response to 20E and the widely studied PLC inhibitor U73122. These findings provide a basis for improved understanding of *A. lucorum* nymph response to 20E, and a more comprehensive understanding of PLC’s function in 20E signal transduction.

## 2. MATERIALS AND METHODS

### Insect Culture

*A. lucorum* specimens were obtained from *Vicia faba* grown in fields in Yancheng (33.110N, 120.250E), Jiangsu Province, China, in July to August 2018. *A. lucorum* nymph and adult individuals underwent rearing on *Phaseolus vulgaris* in a light incubator at 25±1℃ and 70±5% humidity under a 14 h-10 h light/dark cycle.

### Sample Preparation

About 24 h after molting, a total of 600 third instar stage nymphs were collected in plastic cases with green beans. Midguts from early wandering third instar nymphs underwent dissection in Ringer’s solution, and were pre-incubated in 12-well tissue culture plates with 500 μL of Grace’s insect tissue culture medium for 30 min, with or without 50 μM U73211 (Sigma, St. Louis, MO, USA). This was followed by stimulation with 1.0 μM 20E (Sigma, St Louis, MO, USA) or distilled water for 12 h. The assay consisted of four experimental treatments: (1) 20E and U73122; (2) 20E; (3) U73122; (4) CK, negative control, treatment with distilled water. Upon treatment, midguts were obtained, homogenized and submitted to centrifugation (16,000×g, 10 min).

### Protein extraction and Digestion

Specimens were pulverized in liquid nitrogen and treated with 0.5 ml lysis buffer (6 M urea, 2 M thiourea, 4% CHAPS, 40 mM Tris-HCl, 5 mM EDTA, 1 mM PMSF and 10 mM DTT; pH 8.0). After sonication (5 min at 4°C) and centrifugation (18,000×g, 4°C; 15 min), supernatants were collected. Protein amounts were assessed by the Bradford assay. Protein (100 μg) digestion was performed by the FASP method (Tang *et al.*, 2017) utilizing trypsin (Promega, Madison, WI) and 0.1M tetraethylammonium bromide (TEAB) as buffer solution. Two biological replicates were pooled for iTRAQ analysis.

### iTRAQ labeling

After trypsin digestion, lysate specimens (200 μg) were added to 200 μL 8 M urea in 150 mM Tris-HCl (pH 8.0) (buffered urea) and centrifuged (15 min, 14,000×g) in 10 kDa diafiltration tubes for concentration. The concentrated samples were treated as described above for the lysates and incubated at ambient with 100 μL 50 mM iodoacetamide (Bio-Rad, 163-2109, Hercules, CA, USA) in buffered urea and concentrated as described above. After two washes with 100 μL buffered urea by centrifugation (10 min, 14,000×g), the specimens were further resuspended with 100 μL of dissolution buffer (AB SCIEX, Foster City, CA, USA) and centrifuged as above. Next, the pellets in 40 μL dissolution buffer underwent digestion with 2 μg trypsin overnight at 37°C and vacuum drying. Further sample processing with the 8-plex iTRAQ reagent (AB SCIEX) was carried out as directed by the manufacturer. Treated and infected specimens underwent labeling (1 h, ambient) with iTRAQ reagents 113-115 and 116-118, respectively. After vacuum drying, fractionation was carried out by high pH Reversed phase chromatography (Hp-RPC).

### Fractionation by Hp-RPC

A Shimadzu LC-20AB HPLC Pump system was used for Hp-RPC. The iTRAQ-labeled peptide mixture was added to 0.5 mL buffer A (20 mM NH_4_HCO_2_ in 2% acetonitrile, pH 10) followed by injection into a Gemini-NX μM C18 column (110 Å, 250 × 4.6 mm; Phenomenex, Guangzhou, China). Elution was carried out at 1 mL/min initially with buffer A for 10 min, followed by a 5-30% buffer B (20 mM NH_4_HCO_2_ in 98% acetonitrile, pH 10; 15 min) and 30-80% buffer B (3 min) in buffer A. Detection was performed at 214 nm, with fraction collection occurring at 1-min intervals. Finally, the eluates were pooled in 10 fractions and submitted to vacuum drying.

### LC-ESI-MS/MS

Various fractions underwent resuspension in a certain volume of buffer A (5% acetonitrile and 0.1% formic acid in water) and centrifugation (18,000×g, 10 min). The final peptide content was adjusted to approximately 0.5 μg/μl in all fractions. After loading 8 μL of each sample at 3 μL/min on an Eksigent 425 2D HPLC system (AB SCIEX), the peptides were eluted onto an in-housed packed analytical C18 column (internal diameter, 75 μm). A 45-min gradient was initially applied at 0.3 μL/min from 8 to 25% B (84% acetonitrile and 0.1% formic acid), and a 10-min linear gradient to 80% B, which was maintained for 5 min.

A TripleTOF 5600 System was utilized to acquire data, with ion spray voltage at 2.3 kV, curtain and ion source gases at 30 and 6 PSI, respectively, and the chamber at 150°C. MS was carried out in the high-resolution mode (>30,000 fwhm) for TOF MS scanning, with m/z of 350-1800 Da. For information dependent data acquisition (IDA), scan was performed for 50 ms, collecting up to 30 product ions exceeding the cutoff of 200 counts/s and showing a 2+ to 4+ charge.

### iTRAQ protein assessment

The raw data (*.wiff and *.wiff.scan) underwent analysis with ProteinPilot 5.0 (AB SCIEX) against the *A. lucorum* transcriptome database. Search indexes were: Specimen Type, iTRAQ 8plex (Peptide Labeled); Cysteine alkylation, iodoacetamide; Digestion, Trypsin. FDR (false discovery rate) analysis was performed routinely for identification. Differentially expressed proteins (DEPs) were considered with a 2-fold change cutoff.

### Bioinformatics analysis

Two annotation databases, including Gene Ontology (GO) (Ashburner *et al.*, 2000) and Kyoto Encyclopedia of Genes and Genomes (KEGG) (Ogata *et al.*, 1999), were utilized for annotating and grouping DEPs. The GO numbers of DEPs were retrieved with the Blast2GO software (https://www.blast2go.com/), and DEPs were further classified with WEGO 2.0, a web tool for analyzing and plotting GO annotations (Jia *et al.*, 2018). As for KEGG pathway analysis, the protein sequences of DEPs were uploaded onto the KEGG Automatic Annotation Server (Moriya *et al.*, 2007) to compute and group the DEP-integrated signaling pathways.

### Protein analysis

Immunoblot was performed for validating iTRAQ data. In brief, protein were extracted from the midguts post-treatment with Tissue Protein Extraction Reagent Kit (ZoonBio) as directed by the manufacturer. Protein quantitation utilized the bicinchoninic acid (BCA) assay (ZoonBio). Equal amounts of total protein underwent separation by 10% SDS-PAGE and electro-transfer on nitrocellulose membranes (Bio-Rad). After blocking (5% skim milk in Tris-buffered saline with 0.05% Tween 20; 1 h at 37℃), the membranes underwent incubation with primary antibodies targeting trans-sialidase, cuticle protein 19.8, pathogenesis-related protein 5, guanine nucleotide-binding protein and ras-related protein 1 for 1 h at 37℃. Then, horseradish peroxidase (HRP)-linked secondary antibody (ZoonBio) for 1 h at ambient. Development utilized Enhanced Chemiluminescence Advance Kit components (GE Healthcare, PA, USA). Quantitation was carried out by densitometry with ImageJ (http://rsb.info.nih.gov/ij/index.html). Relative protein amounts were obtained by dividing the normalized trans-sialidase, cuticle protein 19.8, pathogenesis-related protein 5, guanine nucleotide-binding protein and ras-related protein 1 densities by the normalized β-actin density based on mouse anti-β-actin antibody (ZoonBio).

## 3. RUSULTS

### Protein profiling and identification of differentially expressed proteins

To obtain a global view of proteome alteration in *A. lucorum* nymphs in response to different stresses (20E, U73122, and 20E+U73122), an iTRAQ-based quantitative proteomic approach was employed to perform comparative analysis among the three groups (20E/control, U73122/control and 20E+U73122/control). A total of 1624 non-redundant proteins with two or more unique peptides were identified (Table S1). Based on the regulation cutoff of 2.0-fold (*p*<0.05), 98 (20 up-regulated and 78 down-regulated), 248 (42 up-regulated and 206 down-regulated), and 266 (57 up-regulated and 209 down-regulated) differentially-expressed proteins (DEPs) were, respectively, identified in the 20E/control, U73122/control and 20E+U73122/control groups (Table S2). For all DEPs among the three groups, the numbers of repressed proteins were higher than those of up-regulated ones, and DEPs were significantly less abundant in the 20E/control group compared with the other groups. Moreover, the numbers of DEPs in the U73122/control and 20E+ U73122/control groups were similar.

Venn diagram analysis (Figure 1, Table S3) further showed that only 8 DEPs were shared among all three groups. From pairwise comparisons of the 20E/control group with the U73122/control and 20E+U73122/control groups, a total of 20 and 18 DEPs were identified, respectively. A total of 155 DEPs were shared between the U73122/control and 20E+U73122/control groups. The numbers of shared proteins were over seven times greater than that of the 20E group. Furthermore, these 155 DEPs exclusively showed the same trend in protein abundance changes (11 were up-regulated and 144 showed down-regulation).

**Figure 1.**
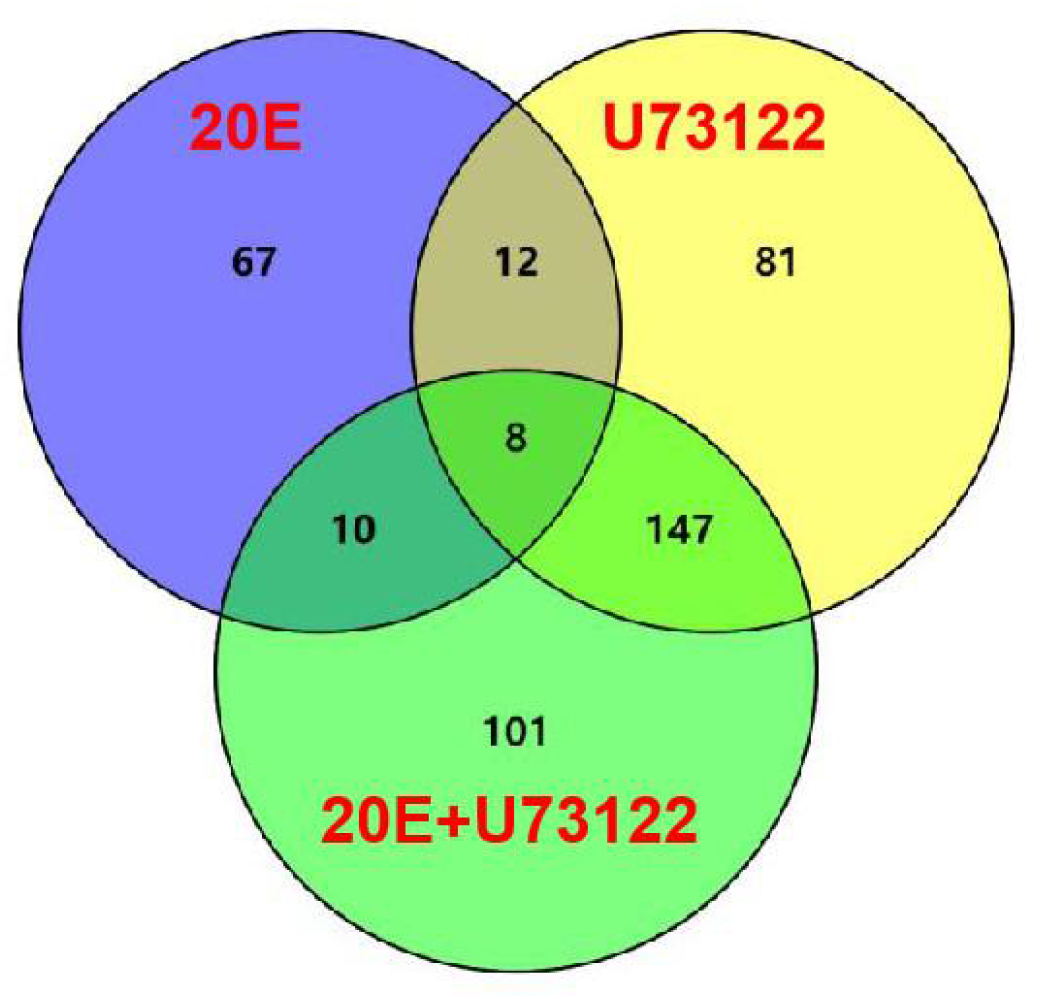
DEPs with a change ≥1.2-fold in *A. lucorum* after treatment with 20E, U73122 and 20E+U73122, respectively. Co-DEPs between two time points and among three time points were also determined.

### Bioinformatics analysis and interpretation of DEPs

Since no complete reference proteome database of *A. lucorum* has been published, we employed the Blast2GO suite (https://www.blast2go.com/), a commercial software, to annotate the DEPs found in each group, with the assigned GO numbers representing their functional relevance in the categories of cellular component (CC), molecular function (MF), and biological process (BP). The GO-assigned DEPs were then classified with WEGO 2.0, a web tool for analyzing and plotting GO annotations (Jia *et al.*, 2018). As shown in Figure 2, more DEPs in the 20E/control group were associated with intracellular localization compared with extracellular distribution. Multiple DEPs possessing catalytic and ion-binding activities were identified in the U73122/control and 20E+U73122/control groups, while many DEPs in the 20E/control group contributed to the regulation of anatomical structure development and stress response.

**Figure 2.**
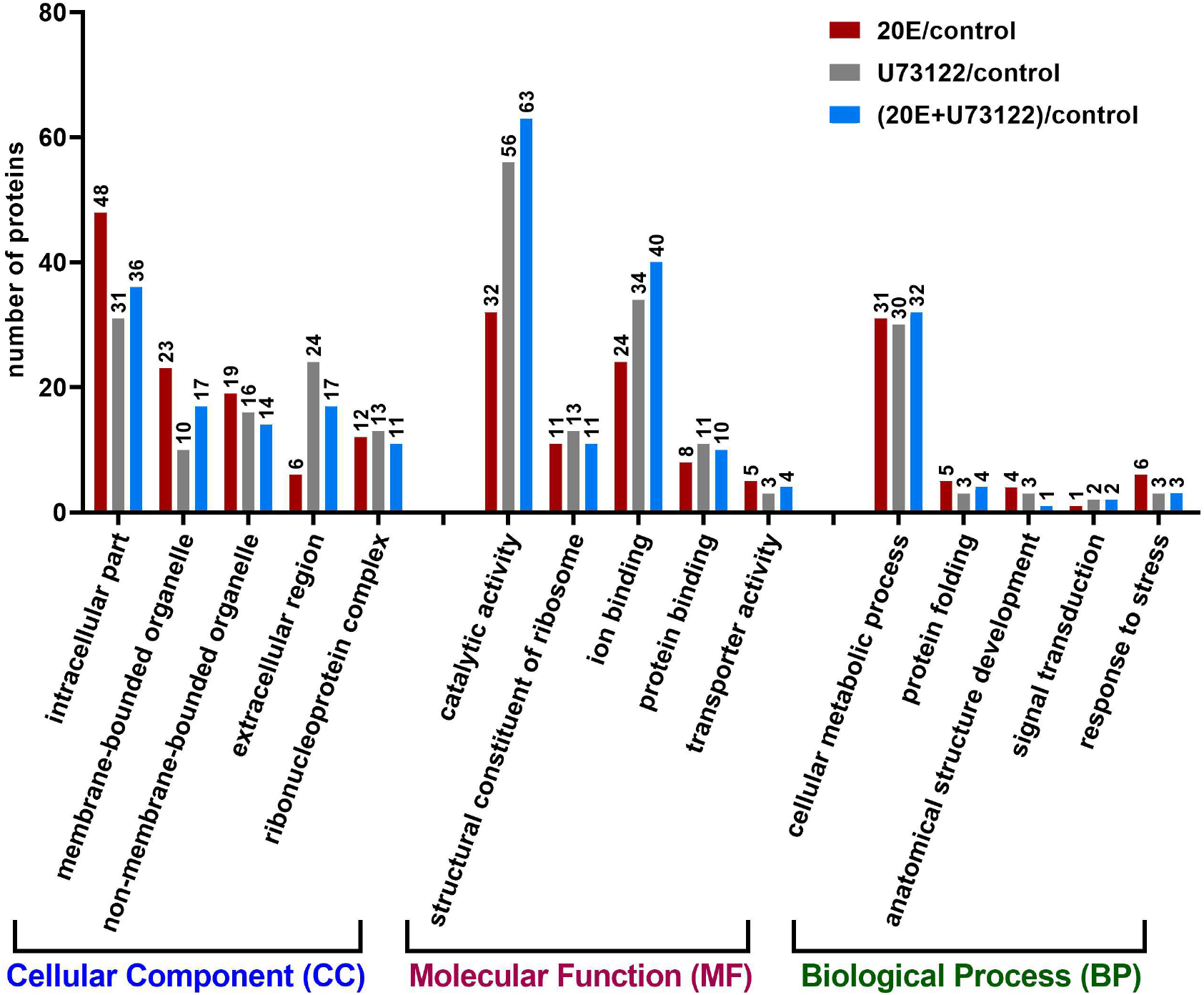
Gene ontology analysis of 612 proteins differentially expressed after treatment with 20E, U73122 and 20E+U73122 of *A. lucorum,* respectively. Proteins were annotated by biological process, cellular component, and molecular function

“Zoomed-in” KEGG pathway analysis revealed that the identified DEPs were integrated into diverse signaling pathways (Figure 3, Table S4). The largest portion of DEPs among the three groups were associated with metabolic pathways. Comparative analysis showed no overlapping DEP among the three groups. All DEPs (n=24) in the 20E/control group were down-regulated, while both up- and down-regulated DEPs were found in the U73122/control (8 up-regulated and 12 down-regulated) and 20E+U73122/control (17 up-regulated and 10 downregulated). Interestingly, multiple ribosomal proteins were identified as DEPs involved in the ribosome process/pathway. In the 20E/control group, all ribosomal proteins (n=11) were down-regulated, while almost all (11 out of totally 12) were up-regulated in the U73122/control group. In the 20E+U73122/control group, the numbers of up- and down-regulated ribosomal proteins were almost even (6:5), suggesting that 20E and U73122 treatments played opposite roles in the regulation of ribosome-driven protein translation.

**Figure 3.**
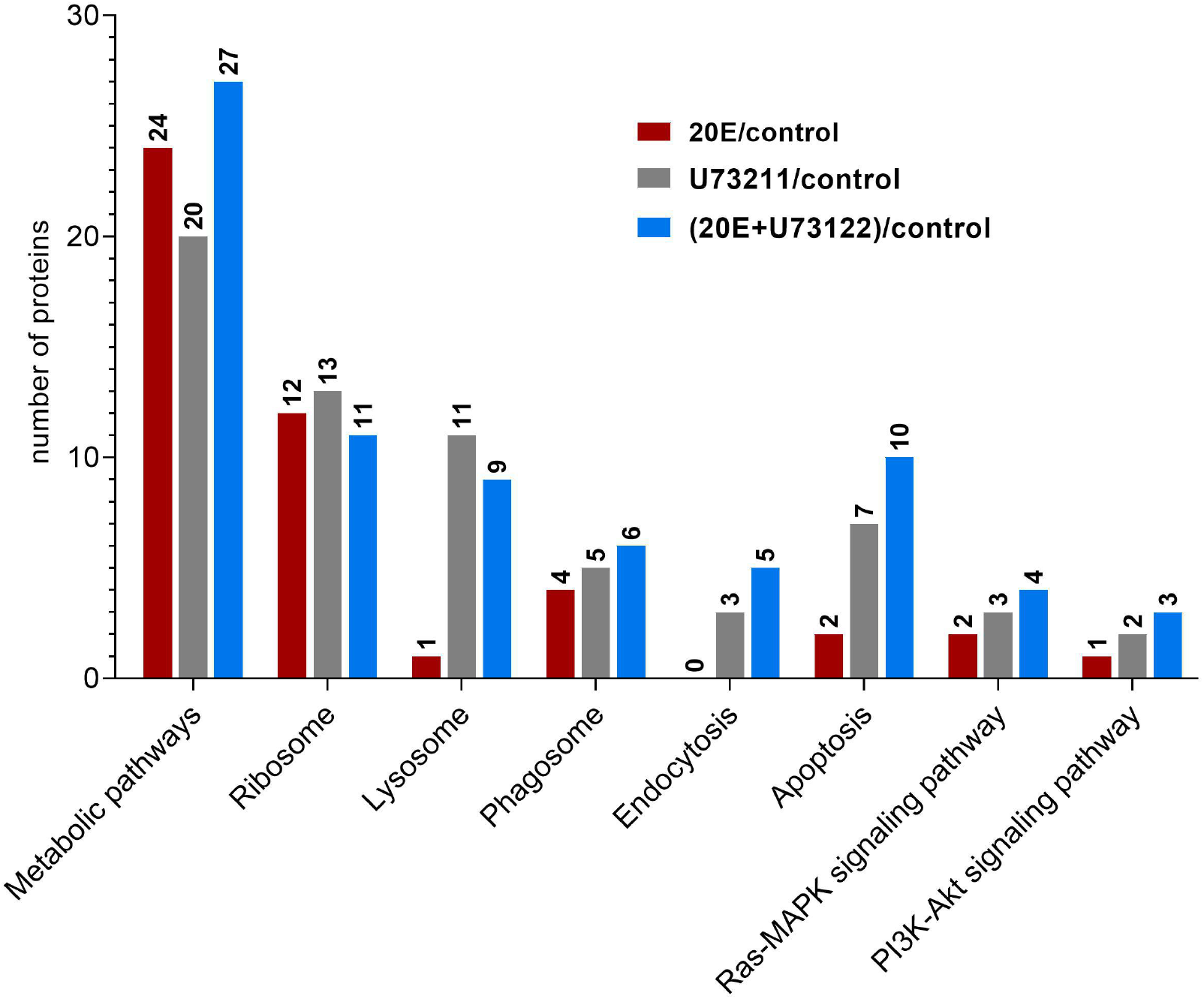
KEGG pathway analysis of DEPs identified after treatment with 20E, U73122 and 20E+U73122 of *A. lucorum*.

Compared with the 20E/control group, DEPs in the U73122/control and 20E+U73122/control groups were more enriched in the processes/pathways of lysosome, phagosome, endocytosis, phagosome (phagocytosis), endosome (endocytosis) and lysosome. A total of 20 non-redundant DEPs in the U73122/control and 20E+U73122/control groups, including 11 shared ones, were involved in these processes. Almost all these DEPs were down-regulated (16 out of 20). In contrast, only 4 DEPs (1 up-regulated and 3 down-regulated) in the 20E/control group were involved in these processes/pathways. Furthermore, several subunits (A, C, G, and E) composed of V-type proton ATPase (ATPeV) were identified as DEPs and classified into the phagosome group. Subunits A (catalytic unit) and G, identified in the 20E+U73122/control and U73122/control groups, respectively, were up-regulated, while subunits C and E (assembling the peripheral stalk to insert the complex into the membrane) were down-regulated in the 20E/control group.

### Validation of DEPs

Western blot showed that trans-sialidase, cuticle protein, and pathogenesis-related protein were clearly up-regulated by 20E, and down-regulated upon treatment with U73122 and 20E+U73122, respectively. Ras-related protein 1 was obviously up-regulated (Figure 4) in response to U73122 and 20E+U73122 treatments but clearly down-regulated by 20E. Guanine nucleotide-binding protein was clearly up-regulated by 20E+U73122, while treatment with 20E and U73122 showed no significant effects on its expression. Results of Western blot confirmed and highlighted the above proteomic findings.

**Figure 4.**
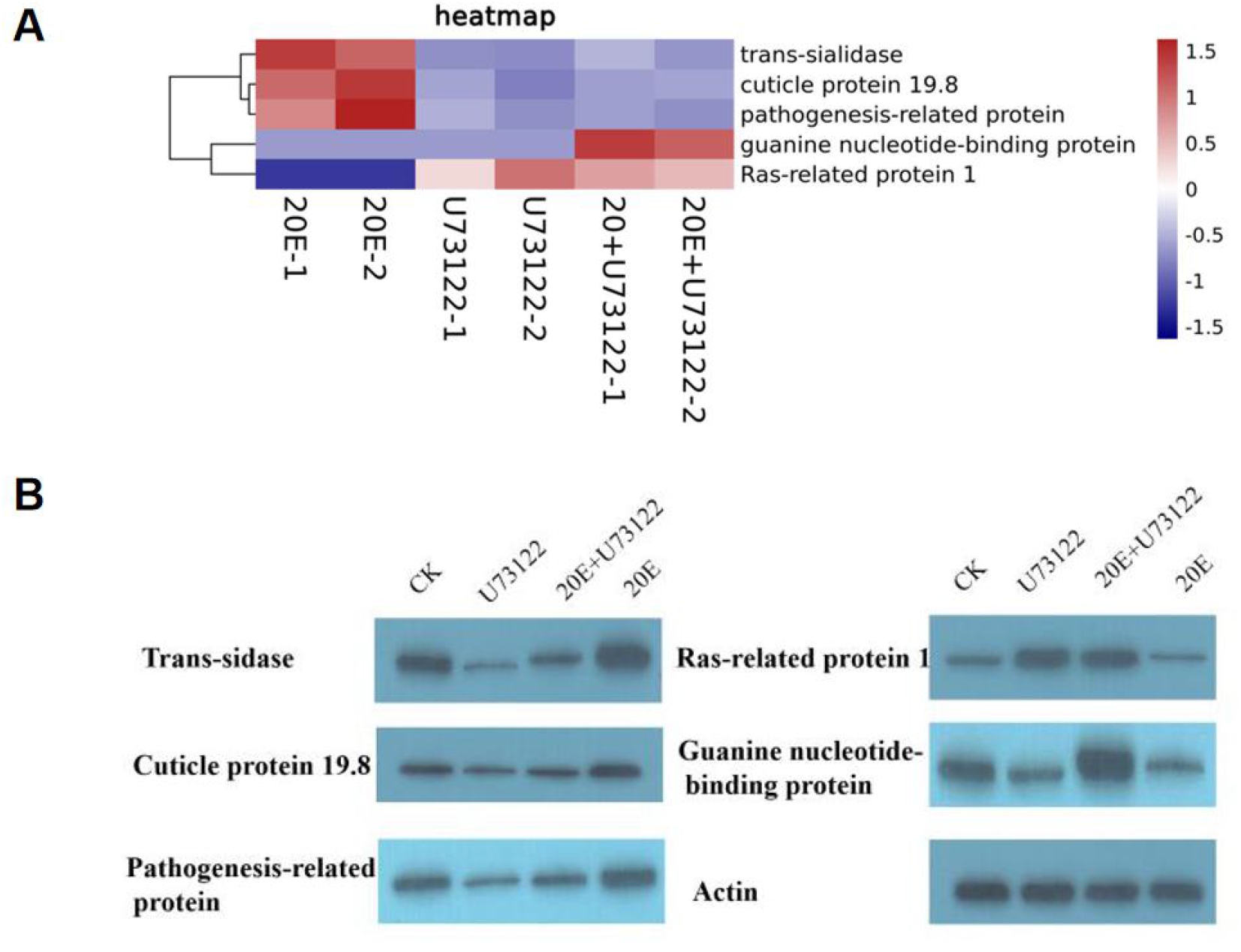
(A) Heat map of Ras-related protein 1 (Rap1), cuticle protein 19.8, guanine nucleotide-binding protein, trans-sialidase and pathogenesis-related protein in different treatment groups of *A. lucorum* in ITRAQ analysis. (B) Western blot of Ras-related protein 1 (Rap1), cuticle protein 19.8, guanine nucleotide-binding protein, trans-sialidase and pathogenesis-related protein in different treatment groups of *A. lucorum*. *Actin* was used for normalization.

## 4. DISCUSSION

Two-dimensional polyacrylamide gel electrophoresis (2-DE), combined with MS, peptide sequencing and database search, represents a great tool for determining DEPs, but is relatively complex and unsuitable for quantitating proteins in minute quantities. iTRAQ, a novel stable isotope technique for protein measurement that employs mass spectrometry (Zieske *et al.*, 2006; Pierce *et al.*, 2008), could comparatively assess eight distinct specimens concurrently and examine proteome profiles upon treatment with exogenous substances.

Ecdysteroids represent a group of steroid hormones regulating developmental, reproductive and other critical biological events in insects. Meanwhile, 20E is considered the major ecdysteroid in most insects (Spindler-Barth *et al.*, 2000; Lafont *et al.*, 2005). Previous evidence indicates that 20E exerts effects through its receptor (Thomas *et al.*, 1993; Hill *et al.*, 2013; Levin, 2015), a heterodimer comprising two nuclear receptors, i.e., ecdysteroid receptor (EcR) and ultraspiracle (Usp), an ortholog of the retinoid X receptor (RXR) in vertebrates (Yao *et al.*, 1992). In this study, after treatment with 20E of *A. lucorum* nymphs, 98 (20 up-regulated and 78 down-regulated) DEPs were identified, and 248 (42 up-regulated and 206 down-regulated) and 266 (57 up-regulated and 209 down-regulated) were, respectively, identified in the U73122/control and 20E+U73122/control groups. Venn diagram analysis further showed that only 8 DEPs were shared in all three groups. Four proteins, including pathogenesis-related protein 5-like, cuticle protein 19.8, trans-sialidase, and larval cuticle protein A2B-like, were up-regulated after 20E treatment but down-regulated in both the U73122/control and 20E+U73122/control groups. Meanwhile, cathepsin L1, protein takeout (hemolymph juvenile hormone binding protein), ATP-dependent RNA helicase p62-like, and myosin-9 isoform X1 were all down-regulated. Pairwise comparisons of the 20E/control group with U73122/control and 20E+U73122/control groups, respectively, revealed 20 and 18 DEPs, indicating that the panels of proteins regulated by 20E and PLC were significantly different, in agreement with the nature of these two reagents evoking distinct cellular responses (Liu **et al.**, 2014; Chen *et al.*, 2019; Tan *et al.*, 2020).

The cuticle of insects has multiple layers and three functional regions, including the epicuticle, procuticle and endocuticle, differing in protein profile, structural properties and physiological roles (Willis *et al.*, 1996). The properties of insect cuticle depend on the construction of cuticular proteins, and various cuticle types show pronounced differences in mechanical features in association with the characteristics of individual proteins (Andersen, 2011). As shown above, all identified cuticular proteins were up-regulated to varying degrees after 20E treatment (Table 1). Examples include larval cuticle protein A2B-like, pupal cuticle protein C1B-like, cuticle protein 19.8 and cuticle protein 19-like. This indicated that cuticular proteins are induced by ecdysone addition in *A. lucorum* and 20E could evoke the up-regulation of these cuticular proteins. Insects produce new cuticles each time the larva molts and various cuticle types are required when the larva changes to the pupa, or when the latter metamorphoses into the adult animal. Mounting evidence suggests that ecdysone signaling is critical for cuticle biosynthesis and accumulation (Riddiford *et al.*, 2001; Kozlova and Thummel, 2003). The production of cuticular proteins via the ecdysone and juvenile hormones is well-known in multiple insects (Charles, 2010). In *Bombyx mori*, several classes of cuticular genes with distinct profiles during development as well as various ecdysone responses from wing disc have been reported (Zhong *et al.*, 2006; Futahashi *et al.*, 2008). Moreover, the temporal expression of CPR95 is regulated by the ecdysone-responsive transcription factor E74A in the wing discs of pre-pupa (Wang *et al.*, 2010). Indeed, most cuticular genes are induced by an ecdysteroid pulse, with their expression requiring the existence and suppression of 20E (Nita *et al.*, 2009; Soares *et al.*, 2007; Wang *et al.*, 2009; Zhong *et al.*, 2006). This is similar to what reported at the stage around ecdysis (Lemoine *et al.*, 2004; Noji *et al.*, 2003; Wang *et al.*, 2009). Remarkably, 12 cuticular proteins and their homologous molecules were down-regulated after U73122 treatment, while 7 cuticular proteins and their homologous molecules were down-regulated by 20E+U73122 (Table 2 and 3). Phospholipase C represents a group of enzymes contributing to the cellular turnover of inositol-containing phospholipids (Dai *et al.*, 2015). The latter enzymes remove the polar head group from membrane lipids, including phosphatidylinositol 4,5-bisphosphate (PIP_2_), for generating inositol 1,4,5-triphosphate and diacylglycerol, which are secondary messengers mobilizing calcium in the cell and activating protein kinase C, separately (Klein *et al.*, 2011). It has been reported that PLC is involved in 20E signal transduction. In this study, while PLC activity was inhibited by the addition of exogenous U733122, it caused suppression of cuticular proteins and their homologous molecules, consistent with Western blot results. Cuticle protein 19.8 was clearly upregulated by 20E, but downregulated upon treatment with U73122 and 20+U73122. This indicates that PLC may participate in the cuticle protein biosynthesis via regulation by ecdysone.

**Table 1.**
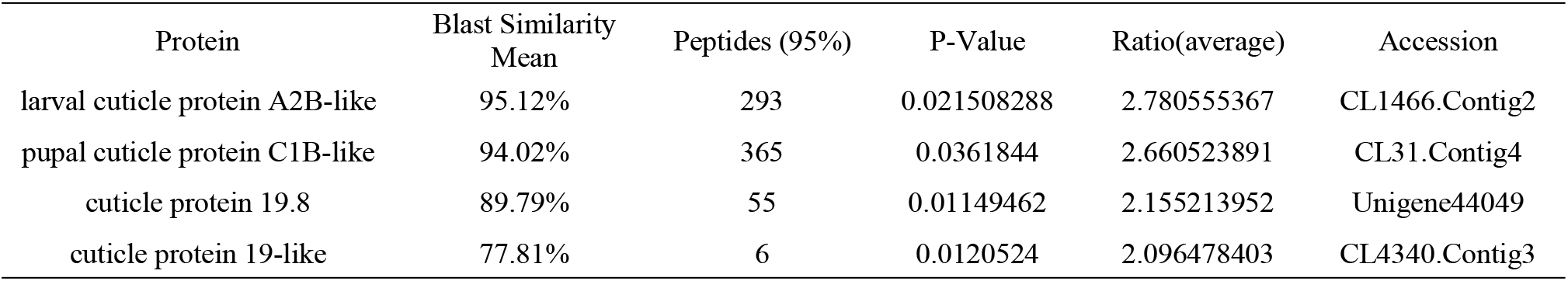
Cuticle proteins and homologies in 20E-treated *A. lucorum*

**Table 2.**
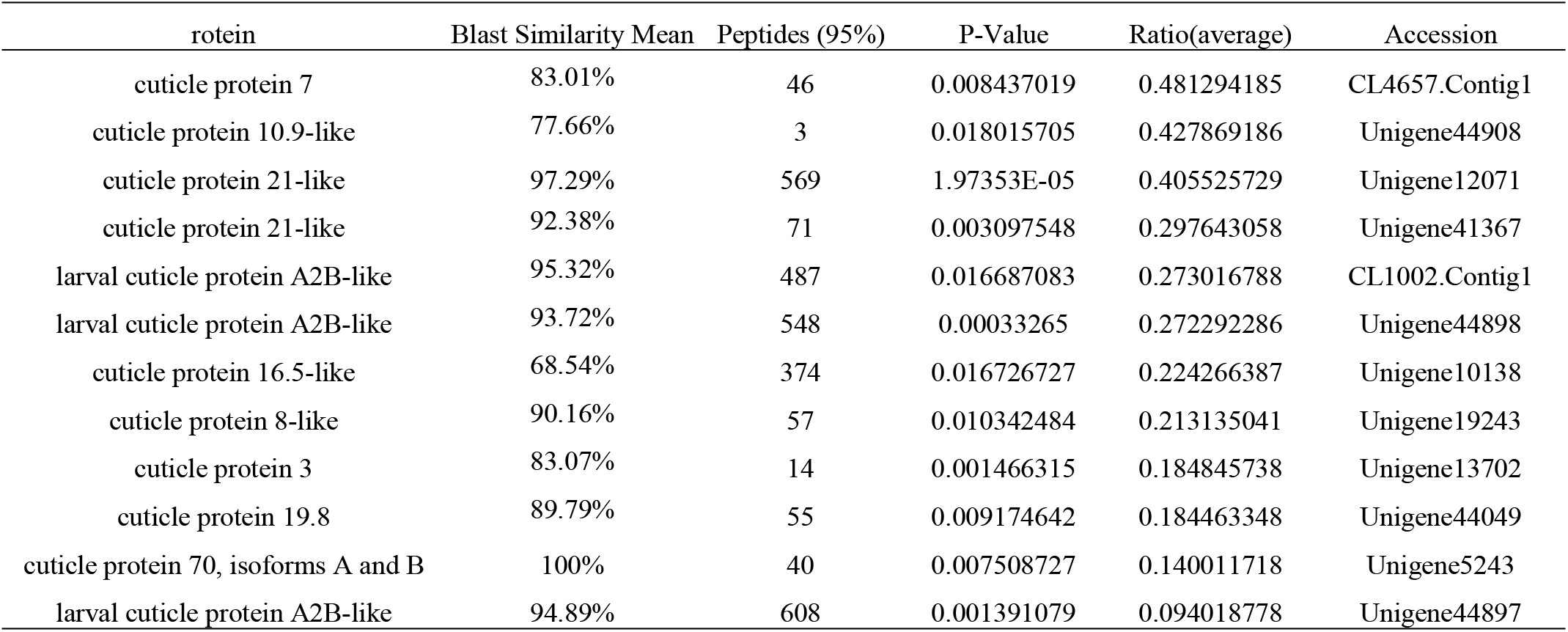
Cuticle proteins and homologies in U73122-treated *A. lucorum*

| rotein | Blast Similarity Mean | Peptides (95%) | P-Value | Ratio(average) | Accession |
| --- | --- | --- | --- | --- | --- |
| cuticle protein 7 | 83.01% | 46 | 0.008437019 | 0.481294185 | CL4657.Contig1 |
| cuticle protein 10.9-like | 77.66% | 3 | 0.018015705 | 0.427869186 | Unigene44908 |
| cuticle protein 21-like | 97.29% | 569 | 1.97353E-05 | 0.405525729 | Unigene12071 |
| cuticle protein 21-like | 92.38% | 71 | 0.003097548 | 0.297643058 | Unigene41367 |
| larval cuticle protein A2B-like | 95.32% | 487 | 0.016687083 | 0.273016788 | CL1002.Contig1 |
| larval cuticle protein A2B-like | 93.72% | 548 | 0.00033265 | 0.272292286 | Unigene44898 |
| cuticle protein 16.5-like | 68.54% | 374 | 0.016726727 | 0.224266387 | Unigene10138 |
| cuticle protein 8-like | 90.16% | 57 | 0.010342484 | 0.213135041 | Unigene19243 |
| cuticle protein 3 | 83.07% | 14 | 0.001466315 | 0.184845738 | Unigene13702 |
| cuticle protein 19.8 | 89.79% | 55 | 0.009174642 | 0.184463348 | Unigene44049 |
| cuticle protein 70, isoforms A and B | 100% | 40 | 0.007508727 | 0.140011718 | Unigene5243 |
| larval cuticle protein A2B-like | 94.89% | 608 | 0.001391079 | 0.094018778 | Unigene44897 |

**Table 3.**
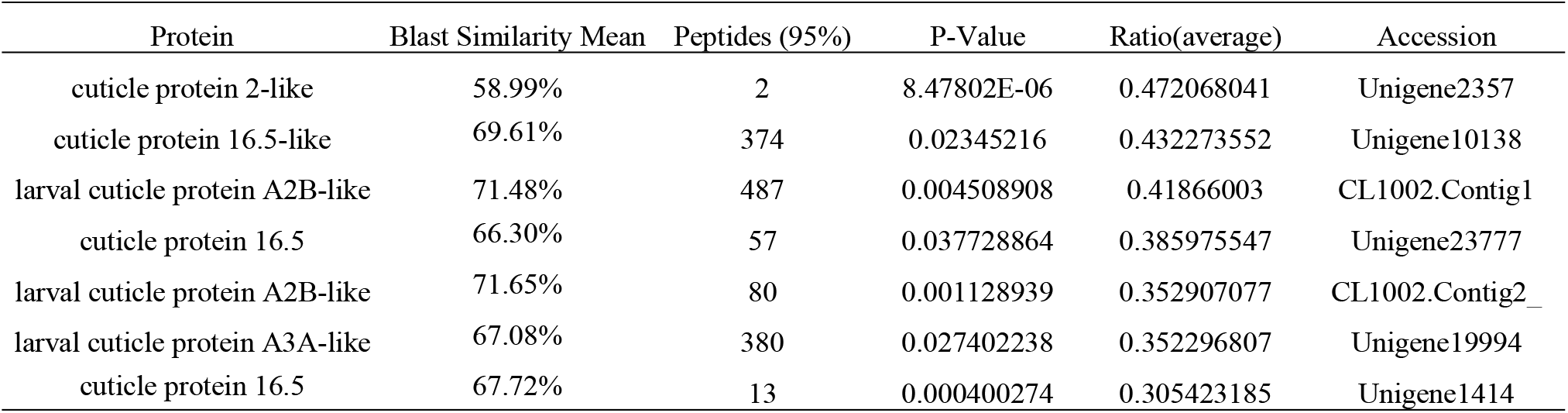
Cuticle proteins and homologies in 20+U73122-treated *A. lucorum*

Besides affecting development, molting, metamorphosis and reproduction, 20E is also involved in the regulation of insect innate immunity. Previous reports demonstrated that 20E induces apoptosis in insect metamorphosis and pathologies. In *B. mori* Bm-12 cells, 20E induces cell cytotoxicity sequentially via autophagy and apoptosis (Xie *et al.*, 2016). The induction and repression of immune reactions by 20E are species- and developmental stage-specific. 20E upregulates prophenoloxidase 1 in *Anopheles gambiae* (Ahmed *et al.*, 1999; Muller *et al.*, 1999) as well as hemolin in the fat bodies of cecropia moth at the diapausing stage (Roxstrom *et al.*, 2005). As shown above, 2, 7 and 10 DEPs were found in the 20E/control, U73122/control and 20E+U73122/control groups, respectively, in the apoptosis pathway. Almost all DEPs among the three groups were down-regulated except spectrin alpha (SPTA) and tubulin alpha-1 in the 20E+U73122/control group. SPTA, a cytoskeleton with binding sites for several proteins, is cleaved during apoptosis to alter membrane stability and form apoptotic bodies. Up-regulation of SPTA by co-treatment of *A. lucorum* with 20E and U73122 may promote cell survival. Jointly, the above data indicate that 20E and U73211 negatively regulate apoptosis, and 20E in combination with U73122 may perform a double-edged role in this pathway. Besides positive effects, 20E also suppresses immune response as reported in *Calliphora vicina*, *Bombyx mori* and *Drosophila*, with downregulated AMPs under high ecdysone amounts at the pupal stage, indicating ecdysone blocks innate immunity (Chernysh *et al.*, 1995; Beckstead *et al.*, 2005; Tian *et al.*, 2010). In the larval molt and during larva to pupa transformation, injected ecdysone induces the down-regulation of AMPs, including lebocin-3, gloverin-like protein 2, and nuecin, in the fat bodies of *B. mori* (Tian *et al.*, 2010). The above discrepant findings indicate that complex mechanisms govern ecdysone’s function in innate immunity.

On the other hand, DEPs in the U73122/control and 20E+U73122/control groups were more enriched in the processes/pathways of lysosome, phagosome, and endocytosis. Phagosome (phagocytosis), endosome (endocytosis), and lysosome are inter-crossed organelles for the degradation and recycling of macromolecules (Braun and Niedergang, 2006). A total of 20 non-redundant DEPs in the U73122/control and (20E+U73122)/control groups, including 11 shared ones, were involved in these processes, and almost all these DEPs were down-regulated (16 out of 20). In contrast, only 4 DEPs (1 up-regulated and 3 down-regulated) in the 20E/control group participated in these processes/pathways. These data indicate that PLC predominantly regulates the processes/pathways of phagosome, endosome, and lysosome in a negative manner. PLC is known to contribute to the differentiation/activation of cells controlling both innate and adaptive branches of the immune system. PLC also affects tyrosine kinase-associated pathways for inhibiting Stat5 via recruitment of the protein-tyrosine phosphatase SHP-1, with which PLC and Stat5 interact in the SPS complex. This complex substantially regulates immune cell activation (Kawakami and Xiao, 2014). Furthermore, several subunits (A, C, G and E) composed of V-type proton ATPase (ATPeV) have been identified as DEPs classified into the phagosome group. ATPeV is a multi-subunit protein of up to 14 different polypeptides located on the organelle membrane. It is involved in the acidification of endocytic and phagocytic organelles to regulate ligand release, macromolecule breakdown, and cation uptake. Subunits A (catalytic unit) and G, identified in the 20E+U73122/control and U73122/control groups, respectively, were up-regulated, while subunits C and E (assembling the peripheral stalk to insert the complex into the membrane) were down-regulated in the 20E/control group, indicating that 20E and U73122 have opposite functions in the regulation of ATPeV activity and the subsequent phagosome process/pathway.

The DEPs among the three groups were also found to regulate the Ras-MAPK and PI3K-AKT pathways. In response to stimuli, activated Ras, a small G protein, triggers the MAPK (RAF-MEK-ERK) and PI3K-AKT cascades to control critical cell events, including growth, proliferation, differentiation, migration and apoptosis (Bonni *et al.*, 1999; Zhang *et al.*, 2014). Ras-related protein 1 (Rap1) was identified as a DEP in both the U73122/control and 20E+U73122/control groups with increased expression, consistent with Western blot results indicating that Rap1 was clearly up-regulated by U73122. Rap1 was reported to interfere with mitogen-activated protein kinase (MAPK) signaling by trapping the RAF protein. In insects, ecdysteroidogenesis by prothoracic glands (PGs) is controlled by the brain peptide prothoracicotropic hormone (PTTH) (Smith and Rybczynski, 2012; Johnson *et al.*, 2013). The mature PTTH interacts with and induces its receptor Torso, a *D. melanogaster* receptor tyrosine kinase (Grillo *et al.*, 2012; Johnson *et al.*, 2013). Induced Torso activates MAPK signaling that encompasses Rap, RAF, MAPK kinase (MEK) and extracellular signal-regulated kinase (ERK), stimulating ecdysteroidogenesis. External U73122 suppressed the expression of Rap1, suggesting that PLC negatively regulates the Ras-RAF-MEK-ERK pathway, which is involved in the biosynthesis of 20E. Furthermore, guanine nucleotide-binding protein G(I)/G(S)/G(T) subunit beta-1 (GNB1) was found to be an up-regulated DEP in the 20E+U73122/control group, and Western blot confirmed these results. GNB1 is a subunit of the heterotrimeric G protein. Its main function is to form a G-beta dimer with the G protein γ subunit (Gγ), and to generate a heterotrimeric G protein with G protein α subunit (Gα). Heterotrimeric G protein could receive the signal transferred by GPCR and pass it to downstream molecules. It has been reported that G protein-coupled receptors (GPCRs) contribute to 20E as a membrane receptor in signaling pathways and transfer the signals into the intracellular space to open calcium channels, causing programmed cell death (Oldham and Hamm, 2007). Interestingly, PLC could be stimulated by GNB1 (Camps *et al.*, 1992). In the lepidopteran insect *H. armigera*, 20E regulates calponin phosphorylation and translocation into the nucleus (Liu *et al.*, 2011). 20E increases intracellular calcium levels via the ecdysone-associated GPCR (Cai *et al.*, 2014), and 20E through G-protein induction increases intracellular Ca^2+^ (Ren *et al.*, 2014). 20E controls gene transcription via the GPCR, G-protein, PLC, calcium, and PKC non-genomic pathway (Liu *et al.*, 2014). Another ecdysone responsible GPCR, termed ErGPCR2, also contributes to 20E-induced non-genomic biological events in *H. armigera* (Jing *et al.*, 2015), indicating that 20E cooperates with U73122 to positively regulate GPCR-mediated non-genomic pathway. In contrast, 14-3-3ζ with down-regulated expression was identified as a DEP in the 20E/control group. In addition, 14-3-3ζ, the adaptor protein, interacts directly with RAF to serve as a negative regulator of RAF function. Furthermore, 14-3-3ζ also activates PI3K via binding of the p85 regulatory subunit, suggesting that 20E exerts different roles in the Ras-MAPK and PI3K-AKT pathways. Taken together, these data showed that 20E and U73122 exert opposite roles in Ras-MAPK and PI3K-AKT pathway regulation.

In summary, 98 (20 up-regulated and 78 down-regulated), 248 (42 up-regulated and 206 down-regulated) and 266 (57 up-regulated and 209 down-regulated) DEPs were differentially expressed in the 20E/control, U73122/control and (20E+U73122)/control groups. The functions of these proteins in 20E signaling as well as responses to 20E deserve further attention. For example, cuticular proteins were overtly involved. In addition, 20E was shown to modulate immune response in *A. lucorum*. Moreover, these findings provide a more comprehensive understanding of PLC’s function in 20E signal transduction and protein interactions involving PLC under 20E regulation.

## Acknowledgements

This study was supported by the Chinese Agricultural Research System (CARS-15-18), the Integration Research and Demonstration of the Technology of Cotton Fertilizer and Pesticide Reduction of China (2017YFD0201900), Jiangsu Agricultural Science and Technology Innovation Fund (CX(19)3098), the National Natural Science Foundation of China (31301668) and JAAS Research Foundation (6111613).

## References

Ahmed, A., Martin, D., Manetti, A.G.O., Han, S.J., Lee, W.J.M. Genomic structure and ecdysone regulation of the prophenoloxidase 1 gene in the malaria vector Anopheles gambiae. (1999). Proc Natl Acad Sci U.S.A. 96, 14795–14800.

Andersen, S. O. Are structural proteins in insect cuticles dominated by intrinsically disordered regions? (2011). Insect Biochem Mol Biol. 41(8), 620–627.

Ashburner, M., Ball, C.A., Blake, J.A., Botstein, D., Butler, H., Cherry, J.M., Davis, A.P., Dolinski, K., Dwight, S.S., Eppig, J.T., Harris, M. A., Hill, D. P., Issel-Tarver, L., Kasarskis, A., Lewis,S., Matese,J. C., Richardson,J. E., Ringwald,M.,Rubin,G. M.,Sherlock,G. Gene ontology: tool for the unification of biology. The Gene Ontology Consortium. (2000). Nat Genet. 25(1), 25–29.

Beckstead, R.B., Lam, G., Thummel, C.S. The genomic response to 20-hydroxyecdysone at the onset of *Drosophila* metamorphosis. (2005). Genome Biol. 6, 1866–1870.

Bonni, A., Brunet, A., West, A. E., Datta, S. R., Takasu, M. A., Greenberg, M. E., Cell survival promoted by the Ras-MAPK signaling pathway by transcription-dependent and -independent mechanisms. (1999). Science. 286(5443), 1358–

Boulanger, A., Dura, J.M. Nuclear receptors and *Drosophila* neuronal remodeling.(2015). Biochim Biophys Acta. 1849, 187–195.

Braun, V., Niedergang, F. Linking exocytosis and endocytosis during phagocytosis. (2012). Biol Cell. 98(3), 195–201.

Cai, M.J., Dong, D.J., Wang, Y., Liu, P.C., Liu, W., Wang, J.X., Zhao, X.F. G-protein-coupled receptor participates in 20-hydroxyecdysone signaling on the plasma membrane. (2014). Cell Commun Signaling. 10.1186/1478-811X-12-9.

Camps, M., Hou, C., Sidiropoulos, D., Stock, J.B., Jakobs, K.H., Gierschik, P. Stimulation of phospholipase C by guanine-nucleotide-binding protein βγ subunits. (1992). Febs Journal. 206(3), 821–831.

Caubet, C., Lacroix, S., Decramer, J., Drube, J.H., Ehrich, H., Mischak, J.L., Bascands, J.P. Schanstra. Advances in urinary proteome analysis and biomarker discovery in pediatric renal disease.(2010). Pediatr Nephrol. 25, 27–35.

Charles, J.P. The regulation of expression of insect cuticle protein genes. Insect Biochem Mol Biol. (2010). 40(3), 205–213.

Chen, C.H., Di, Y.Q., Shen, Q.Y., Wang, J.X., Zhao, X.F. The steroid hormone 20-hydroxyecdysone induces phosphorylation and aggregation of stromal interacting molecule 1 for store-operated calcium entry. (2019). J Biol Chem. 294(41), 14922–14936 .

Chernysh, S.I., Simonenko, N.P., Braun, A., Meister, M. Developmental variability of the antibacterial response in larvae and pupae of *Calliphora Vicina* (Diptera, Calliphoridae) and Drosophila melanogaster (Diptera Drosophilidae). (1995). Eur J Entomol. 92, 203–209.

Christiaens, O., Iga, M.,Velarde, R.A., Rouge, P., Smagghe, G. Halloween genes and nuclear receptors in ecdysteroid biosynthesis and signalling in the pea aphid. (2010). Insect Mol Biol. 19,187–200.

Dai, L., Zhuang, L.H., Zhang, B.C., Wang, F., Chen, X.L., Xia, C. Dag/pkcδ and ip3/ca2+/camk iiβ operate in parallel to each other in plcγ1-driven cell proliferation and migration of human gastric adenocarcinoma cells, through akt/mtor/s6 pathway. (2015). Int J Mol Sci, 16, 28510–28522.

Falkenstein, E., Tillmann, H. C., Christ, M., Feuring, M., Wehling, M. Multiple actions of steroid hormones: a focus on rapid, nongenomic effects. (2000). Pharmacol Rev. 52, 513–556

Futahashi, R., Okamoto, S., Kawasaki, H., Zhong, Y.S., Iwanaga, M., Mita, K., Fujiwara, H. Genome-wide identification of cuticular protein genes in the silkworm, *Bombyx mori*.(2008). Insect Biochem. Mol. Biol. 38, 1138–1146.

Grillo, M., Furriols, M., de Miguel, C., Franch-Marro, X., Casanova, J. Conserved and divergent elements in torso RTK activation in *Drosophila* development. (2012). Sci. Rep. 2, 2061–2072.

Hill, R.J., Billas, I.M., Bonneton, F., Graham, L.D., Lawrence, M.C. Ecdysone receptors: from the Ashburner model to structural biology. (2013). Annu Rev Entomol. 58, 251–71.

Iga, M., Iwami, M., Sakurai, S. Nongenomic action of an insect steroid hormone in steroid-induced programmed cell death. (2007). Mol Cell Endocrinol. 263, 18–28.

Iwema, T., Billas, I., Beck, Y., Bonneton, F., Nierengarten, H., Chaumot, A., Richards, G., Laudet, V., Moras, D. Structural and functional characterization of a novel type of ligand-independent RXR-USP receptor. (2007). EMBO J. 26, 3770–3782.

Jing, Y.P., Liu, W., Wang J.X., Zhao, X.F. The steroid hormone 20-hydroxyecdysone via nongenomic pathway activates Ca^2+^/calmodulin-dependent protein kinase II to regulate gene expression. (2015). J Biol Chem. 290(13), 8469–8481.

Johnson, T.K., Crossman, T., Foote, K.A., Henstridge, M.A., Saligari, M.J., Forbes Beadle, L.,Herr, A., Whisstock, J.C., Warr, C.G. Torso-like functions independently of torso to regulate *Drosophila* growth and developmental timing. (2013). Proc Natl Acad Sci U. S. A. 110, 14688–14692.

Kawakami, T., Xiao, W. Phospholipase C-β in immune cells. (2013). Adv Biol Regul. 53(3), 249–257.

Klein, R.R., Bourdon, D.M., Costales, C.L., Wagner, C.D., White, W.L.,Williams, J.D., Hicks, S.N., Sondek, J., Thakker, D.R. Direct activation of human phospholipase C by its well known inhibitor U73122. (2011). J Biol Chem. 286(14), 12407–12416.

Kozlova, T., Thummel, C.S. Essential roles for ecdysone signaling during *Drosophila* mid-embryonic development. (2003). Science, 301, 1911–1914.

Lafont, R., Dauphin-Villemant, C., Warren, J.T., Rees, H. (2005). Ecdysteroid hemistry and Biochemistry, Comprehensive Molecular Insect Science. Elsevier, London.

Lemoine, A., Mathelin, L., Braquart-Varnier, C., Everaerts, C., Delachambre, J. A functional analysis of ACP-20, an adult specific cuticular protein gene from the beetle Tenebrio: role of an intronic sequence in transcriptional activation during the late metamorphic period. (2004), Insect Mol Biol. 13, 481–493.

Levin, E.R. Extranuclear steroid receptors are essential for steroid hormone actions. (2015). Annu Rev Med. 66,271–280.

Liu, P. C., Wang, J. X., Song, Q. S., Zhao, X. F. The participation of calponin in the cross talk between 20-hydroxyecdysone and juvenile hormone signaling pathways by phosphorylation variation. (2011). PLoS One. 10.1371/journal.pone.0019776.

Liu, W., Cai, M. J., Wang, J. X., and Zhao, X. F. In a non-genomic action, steroid hormone 20-hydroxyecdysone induces phosphorylation of cyclin-dependent kinase 10 to promote gene transcription. (2014). Endocrinology. 155(5), 1738–1750.

Liu, W., Cai, M. J., Zheng, C. C.,Wang J.X., Zhao, X.F. Phospholipase Cγ1 connects the cell membrane pathway to the nuclear receptor pathway in insect steroid hormone signaling. (2014). J Biol Chem. 289, 13026–13041.

Lu, Y.H., Liang, G.M., Wu, K.M. Advances in integrated management of cotton mirids. (2007). Plant Prot. 33, 10–15.

Lu, Y.H., Wu, K.M. (2008). Biology and control of cotton mirids. Golden Shield Press.

Lu, Y.H., Wu, K.M., Jiang, Y.Y., Xia, B., Li, P., Feng, H.Q. Mirid bug outbreaks in multiple crops correlated with wide-scale adoption of Bt cotton in China. (2010). Science. 328, 1151–1154.

Lu,Y.H., Wu, K.M. Mirid bugs in China: pest status and management strategies.(2011). Outlooks Pest Manage. 22, 248–252.

Manaboon, M., Iga, M., Iwami, M., and Sakurai, S. Intracellular mobilization of Ca^2+^ by the insect steroid hormone 20-hydroxyecdysone during programmed cell death in silkworm anterior silk glands. (2009). J. Insect Physiol. 55, 122–128.

Meldrum, D.R. G-protein-coupled receptor 30 mediates estrogen’s nongenomic effects after hemorrhagic shock and trauma. (2007). Am J Pathol. 170, 1148–1151.

Moriya, Y., Itoh, M., Okuda, S., Yoshizawa, A.C., Kanehisa, M. KAAS: an automatic genome annotation and pathway reconstruction server. (2007). Nucleic Acids Res. 35, 182–185.

Muller, H.M., Dimopoulos, G., Blass, C., Kafatos, F.C. A hemocyte-like cell line established from the malaria vector Anopheles gambiae expresses six prophenoloxidase genes. (1999). J. Biol. Chem. 274, 11727–11735.

Nakagawa, Y., Henrich, V.C. Arthropod nuclear receptors and their role in molting.(2009). FEBS J. 276,6128–6157.

Nita, M., Wang, H.B., Zhong, Y.S., Mita, K., Iwanaga, M., Kawasaki, H. Analysis of ecdysone-pulse responsive region of BMWCP2 in wing disc of *Bombyx mori*. (2009). Comp Biochem Physiol. 153, 101–108.

Noji, T., Ote, M., Takeda, M., Mita, K., Shimada, T., Kawasaki, H. Isolation and comparison of different ecdysone-responsive cuticle protein genes in wing discs of *Bombyx mori*. (2003). Insect Biochem Mol Biol. 33, 671–679.

Ogata, H., Goto, S., Sato, K., Fujibuchi, W., Bono, H., Kanehisa, M. KEGG: Kyoto Encyclopedia of Genes and Genomes. (1999). Nucleic Acids Res. 27(1), 29–34.

Oldham, W.M., Hamm, H.E. Heterotrimeric G protein activation by G-protein-coupled receptors. (2007). Nat Rev Mol Cell Biol. 9(1), 60–71.

Pierce, R.D., Unwin, C.A., Evans, S., Griffiths, L., Carney, L., Zhang, E., Jaworska, C.F., Lee, D., Blinco, M.J., Okoniewski, C.J., Miller, D.A., Bitton, E., Spooncer, A.D. Whetton. Eightchannel iTRAQ enables comparison of the activity of six leukemogenic tyrosine kinases. (2008). Mol Cell Proteomics. 7, 853–863.

Ren, J., Li, X. R., Liu, P. C., Cai, M. J., Liu, W., Wang, J. X., and Zhao, X. F. G-protein q participates in the steroid hormone 20-hydroxyecdysone nongenomic signal transduction. (2014). J. Steroid Biochem. Mol. Biol. 144, 313–323.

Riddiford, L.M., Cherbas, P., Truman, J.W. Ecdysone receptors and their biological actions. (2001). Vitam. Horm. 60, 1–73.

Ross, P.L., Huang, Y.N., Marchese, J.N.,Williamson, B., Parker, K., Hattan, S., Khainovski, Nikita., Pillai, S., Dey, S., Daniels, S., Purkayastha, S., Juhasz, P., Martin, S., Bartlet, J.M., He, F., Jacobson, A., Pappin, D.J. Multiplexed protein quantitation in Saccharomyces cerevisiae using amine-reactive isobaric tagging reagents.(2004). Mol Cell Proteomics. 3, 1154–1169.

Roxstrom, L.K., Assefaw, R.Y., Rosinska, K., Faye, I. 20-Hydroxyecdysone indirectly regulates hemolin expression in *Hyalophora cecropia*. (2005). Insect Mol. Biol. 14, 645–652.

Sandén, C., Broselid, S., Cornmark, L., Andersson, K., Daszkiewicz-Nilsson, J.,Mårtensson, E.A., Olde, B., Leeb-Lundberg, L.M. Fredrik. G Protein-Coupled estrogen receptor 1/G protein-Coupled receptor 30 localizes in the plasma membrane and traffics intracellularly on cytokeratin intermediate filaments. (2011). Mol Pharmacol. 79(3), 400–410.

Smith, T., and Evans, P. D. Rapid, nongenomic responses to ecdysteroids and catecholamines mediated by a novel *Drosophila* G-proteincoupled receptor. (2005). J. Neurosci. 25, 6145–6155

Soares, P.M.M., Elias-Neto, M., Simões, L.P.Z., Bitondi, M.G.M. A cuticle protein gene in the honeybee: expression during development and in relation to the ecdysteroid titer. (2007). Insect Biochem Mol Biol. 37, 1272–1282.

Spindler-Barth, M., Spindeler, K.D. (2000). Hormonal regulation of larval moulting and metamorphosis-Molecular aspects. In: Dorn A (ed) Progress in developmental endocrinology. Wiley-Liss, New York.

Tan, Y.A., Zhao, X.D., Sun, H.J., Zhao, J., Xiao, L.B., Hao, D.J., Jiang, Y.P. Phospholipase C gamma (PLCγ) regulates soluble trehalase in the 20E-induced fecundity of Apolygus lucorum.(2020). Insect science 10.1111/1744-7917.12772.

Thomas, H.E., Stunnenberg, H.G., Stewart, A.F. Heterodimerization of the Drosophila ecdysone receptor with retinoid X receptor and ultraspiracle. (1993). Nature. 362,

Thompson, E. B. Steroid hormones: membrane transporters of steroid hormones. (1995). Curr. Biol. 5, 730–732.

Tian, L., Guo, E., Diao, Y., Zhou, S., Peng, Q. Genome-wide regulation of innate immunity by juvenile hormone and 20-hydroxyecdysone in the Bombyx fat body. (2010). BMC Genomics. 10.1186/1471-2164-11-549.

Wang, H.B., Iwanaga, M., Kawasaki, H. Activation of BMWCP10 promoter and regulation by BR-C Z2 in wing disc of *Bombyx mori*. (2009). Insect Biochem. Mol. Biol. 39, 615–623.

Wang, H.B., Moriyama, M., Iwanaga, M., Kawasaki, H. Ecdysone directly and indirectly regulates a cuticle protein gene, BMWCP10, in the wing disc of Bombyx mori.(2010). Insect Biochem Mol Biol. 40(6), 453–459.

Wang, H.B., Nita, M., Iwanaga, M., Kawasaki, H. βFTZ-F1 and Broad-Complex positively regulate the transcription of the wing cuticle protein gene, BMWCP5, in wing discs of *Bombyx mori*. (2009). Insect Biochem. Mol. Biol. 39, 624–633.

Willis, J.H. Metamorphosis of the Cuticle, Its Proteins, and Their Genes. (1996) Metamorphosis. 208(5), 253–282.

Wu, K.M., Mu, W., Liang, G.M., Guo, Y.Y. Regional reversion of insecticide resistance in *Helicoverpa armigera* (Lepidoptera: Noctuidae) is associated with the use of Bt cotton in northern China. (2005). Pest Manag Sci. 61, 491–498.

Xie, K., Tian, L., Guo, X.Y., Li, K., Li, J.P., Deng, X.J., Li, Q.R., Xia, Q.Y., Zhong Y.J, Huang, Z.J., Liu, J.P., Li, S., Yang, W.Y., Cao, Y. BmATG5 and BmATG6 mediate apoptosis following autophagy induced by 20-hydroxyecdysone or starvation. (2016). Autophagy. 12(2), 381–396.

Xu, T.Q., Jiang, X., Denton, D., Kumar, S. Ecdysone controlled cell and tissue deletion.(2020). Cell Death Differ. 27(1), 1–14.

Yao, T.P., Forman, B.M., Jiang, Z.Y.; Cheabas, L., Chen, J.D., McKeown, M., Cherbas, P., Evans, R.M. Functional ecdysone receptor is the product of EcR and Ultraspiracle genes. (1993). Nature. 366,476–479.

Ye, H., Sun, L., Huang, X., Zhang,P., Zhao, X. A proteomic approach for plasma biomarker discovery with 8-plex iTRAQ labeling and SCX-LC-MS/MS.(2010). Mol. Cell. Biochem. 343, 91–99.

Ye, J., Zhang, Y., Cui, H.H., Liu, J.W., Wu, Y.Q., Cheng, Y., Xu, H.X., Huang, X.X., Li, S.T., Zhou, A., Zhang, X.Q., Bolund, L., Chen Q., Wang J.,Yang, H.M., Fang, L., Shi, C.M. WEGO 2.0: a web tool for analyzing and plotting GO annotations, 2018 update. (2018). Nucleic Acids Research. 46, W1,W71–W75.

Zhang, H., Meng, Q., Tang, P., Li, X., Zhu, W., Zhou, G.L., Shu, R.H., Zhang J.H., Qin, Q.L. Gene expression pattern of insect fat body cells from in vitro challenge to cell line establishment.(2014). In Vitro Cell Dev Biol. 50(10), 952–972.

Zhong, Y.S., Mita, K., Shimada, T., Kawasaki, H. Glycine-rich protein genes,which encode a major component of the cuticle, have different developmental profiles from other cuticle protein genes in *Bombyx mori*. (2006). Insect Biochem Mol Biol. 36, 99–110.

Zieske, L.R. A perspective on the use of iTRAQ reagent technology for protein complex and profiling studies. (2006). J Exp Bot. 57 (7), 1501–1508.

